# Global analysis of a cancer model with drug resistance due to Lamarckian induction and microvesicle transfer

**DOI:** 10.1101/2020.06.22.164392

**Authors:** Attila Dénes, Gergely Röst

## Abstract

Development of resistance to chemotherapy in cancer patients strongly effects the outcome of the treatment. Due to chemotherapeutic agents, resistance can emerge by Darwinian evolution. Besides this, acquired drug resistance may arise via changes in gene expression. A recent discovery in cancer research uncovered a third possibility, indicating that this phenotype conversion can occur through the transfer of microvesicles from resistant to sensitive cells, a mechanism resembling the spread of an infectious agent. We present a model describing the evolution of sensitive and resistant tumour cells considering Darwinian selection, Lamarckian induction and microvesicle transfer. We identify three threshold parameters which determine the existence and stability of the three possible equilibria. Using a simple Dulac function, we give a complete description of the dynamics of the model depending on the three threshold parameters. We demonstrate the possible effects of increasing drug concentration, and characterize the possible bifurcation sequences. Our results show that the presence of microvesicle transfer cannot ruin a therapy that otherwise leads to extinction, however it may doom a partially successful therapy to failure.

## 1 Introduction

Chemotherapy is a method for cancer treatment using anticancer drugs given as a curative agent or with the aim to prolong the patient’s life and reduce the symptoms [1, 16]. During chemotherapy, a single drug or a combination of drugs is usually given at intervals in pulsed doses or cycles. Cytotoxic agents damage tumour cells, which may then lead to cell death, while application of cytostatic drugs suppress tumour growth without direct cytotoxic effect. Chemotherapy resistance – a major difficulty in cancer treatment – means that a tumour previously responsive to the therapy, begins to grow as cancer cells evolve the ability to prevent the development of an effective concentration of the active agent within them. Several ways can lead to resistance of tumour cells to chemotherapy. It has been shown that tumours evolve in a similar way as Darwinian evolution acts, i.e. tumour cells are affected by selective pressure which results in the emergence of the fittest clones [6, 7, 8, 9]. In case of chemotherapy, drugs operate as selective pressure agents. Under their effect, resistant descendant cells arise in the tumour cell population. Another way of development of resistant cells is Lamarckian induction [13], which means that a subpopulation of sensitive cells acquires resistance via changes in gene expression. A third way of appearance of resistance has recently been revealed. It is known that resistance strongly depends on intercellular communication and on tumour microenvironment. Information transfer among tumour and healthy cells affects both local and nonlocal interactions. The latter include long-range cell signalling, delivery of soluble factors and exchange of extracellular vesicles and they are responsible for the active modulation of tumour microenvironment. Microvesicles are extracellular particles released from the cell membrane transporting efflux membrane transporters, genetic information and transcription factors needed for their production in recipient cells. The important role played by microvesicles in the intercellular communication among cancer cells has been revealed by some recent studies [3, 4, 10, 12, 14, 15]. The way how microvesicles emitted by more aggressive donor cells are capable to transport cellular components to less aggressive acceptor cells resembles the transmission of infectious diseases.

In [2], the authors provided experimental evidence from in vitro assays to show that an important exogenous source of resistance is the action of chemotherapeutic agents. This action not only affects the signalling pathways but also the interactions among cells. The authors established a mathematical kinetic transport model consisting of a system of hyperbolic partial differential equations to describe the dynamics displayed by a system of non-small cell lung carcinoma cells exhibiting a complex interplay between Darwinian selection, Lamarckian induction and the nonlocal transfer of extracellular microvesicles. Here we consider a non-spatial version of that system, that allows us to perform a comprehensive mathematical analysis of its dynamics.

To formulate our model, let us denote by *S*(*t*) the number of sensitive cells at time *t* and let *R*(*t*) stand for the number of resistant cells. We denote by *c*(*t*) the drug concentration in the patient’s organism at time *t*. Let *β* denote the rate of microvesicle-mediated transfer from sensitive to resistant cells and *θ* is the cytotoxic action induced cell mortality of sensitive cells due to drugs. The notations *ρ*_0_ and *ρ*_*r*_ stand for reproduction rates of sensitive and resistant cells, respectively, here we assume *ρ*_0_ *> ρ*_*r*_. Parameters *µ*_0_ and *µ*_*r*_ denote death rates of sensitive and resistant cells, respectively, due to apoptosis. For the tumour growth, we assume a logistic form with carrying capacity *K*. The letter *p* stands for the rate of phenotype conversion due to Lamarckian induction. The notations *α* and *ε* stand for drug uptake rate of sensitive and resistant cells, respectively, while *λ*_0_ denotes drug removal rate. The function *I*(*t*) describes time-dependent drug dosage. With these notations, our model takes the form

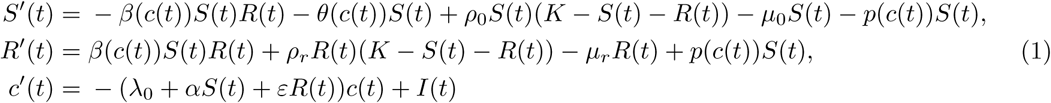

In the next section we make the simplifying assumption that the drug concentration *c*(*t*) is constant, and investigate the case of changing drug concentration later. This assumption transforms (1) into the system

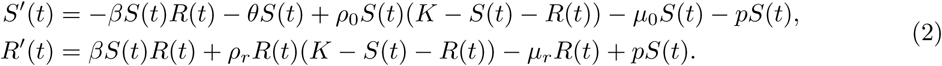

In Section 2, we will give a complete characterization of the global dynamics of system (2) depending on the parameters.

## 2 Description of the global dynamics

### 2.1 Existence and local stability of equilibria

Let us define the following threshold parameters:

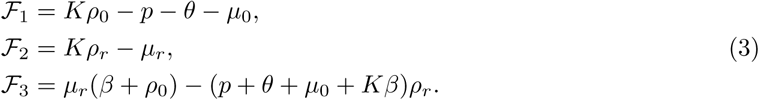

In the following, we will determine the possible equilibria of system (2) and their local stability properties depending on the parameters and describe the dynamics of (2) for all possible combinations of the signs of the above four parameters. To show which combinations of these signs cannot be realized, we prove some simple statements.

Solving the algebraic system of equations

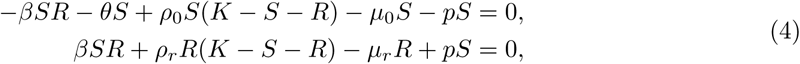

one can easily obtain the trivial equilibrium *E*_0_ = (0, 0) and the equilibrium 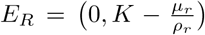 where only resistant cells are present. Because of Lamarckian induction, there is no equilibrium with only sensitive cells. For any coexistence equilibrium (*S*^∗^, *R*^∗^), one obtains that the equality *S*^∗^ = (*ℱ*_1_ − *R*^∗^(*β* + *ρ*_0_))*/ρ*_0_ holds, while *R*^∗^ is given as the solution of the equation *aR*^2^ + *bR* + *c* = 0 with *a* = *β*(*β* + *ρ*_0_ − *ρ*_*r*_), *b* = ℱ_3_ + *β*(ℱ_2_ − ℱ_1_) + *p*(*β* + *ρ*_0_) and *c* = *p*(*p* + *θ* + *µ*_0_ − *Kρ*_0_) = −*p ℱ*_1_. From this, a necessary condition of a coexistence equilibrium is 𝒟:= *b*^2^ − 4*ac >* 0. If, the other way around, we express *R*^∗^ from the first equation, *S*^∗^ is obtained as one of the solutions of the quadratic equation 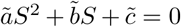 with 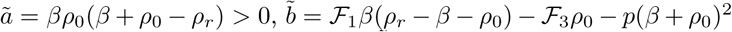 and 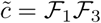. Again, the quadratic equation has real solutions if 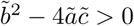, which is equivalent to the condition 𝒟*>* 0.

We prove some simple statements concerning the threshold parameters defined in (3).

#### Proposition 2.1.

*The sensitive cells die out whenever ℱ*_1_ *<* 0.

*Proof*. We estimate *S′*(*t*) as

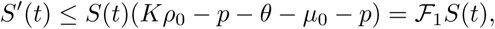

hence, if *ℱ*_1_ *<* 0, then *S*(*t*) → 0 as *t* → ∞. □

#### Remark 2.2.

It follows from Proposition 2.1 that no coexistence equilibrium can exist whenever *ℱ*_1_ *<* 0.

#### Proposition 2.3.

i. ℱ_1_ > 0 *and* ℱ_2_ < 0 *imply* ℱ_3_ > 0.
ii. ℱ_1_ < 0 *and* ℱ_2_ > 0 *imply* ℱ_3_ < 0.
iii. 𝒟< 0 *implies* ℱ_1_ < 0.

*Proof*.

i. Suppose *ℱ*_1_ *>* 0 and *ℱ*_2_ *<* 0 hold but *ℱ*_3_ *<* 0. Then we have *Kρ*_0_ *> p* + *θ* + *µ*_0_ and *Kρ*_*r*_ *< µ*_*r*_, hence, if ℱ_3_ *<* 0 holds, then *µ*_*r*_(*β* + *ρ*_0_) *<* (*p* + *θ* + *µ*_0_)*ρ*_*r*_ *< Kρ*_0_*ρ*_*r*_ *< ρ*_0_*µ*_*r*_, which is a contradiction.
ii. This statement can be shown in an analogous way.
iii. In order to have *𝒟 <* 0, the value −4*βp*(*β* + *ρ*_0_ − *ρ*_*r*_) *ℱ*_1_ has to be positive, which can only happen is *ℱ*_1_ *<* 0.

The following simple statement concerning the existence of a coexistence equilibrium will be useful during the complete description of the global dynamics of system (2).

#### Proposition 2.4.

*If ℱ*_1_ *>* 0 *and ℱ*_3_ *<* 0 *then there is no coexistence equilibrium*.

*Proof*. If *𝒟<* 0 then there is no coexistence equilibrium. Let us suppose that *𝒟>* 0. From Vieta’s formulas, we obtain that if ℱ_1_ *>* 0 and ℱ_3_ *<* 0 then the equation *aR*^2^ + *bR* + *c* = 0 has exactly one positive solution as *a >* 0 and *c/a <* 0. Similarly, the equation 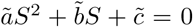 also has exactly one positive solution as *ã* > 0 and 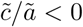. However, we now that for any coexistence equilibrium, the equality *ρ*_0_*S*^∗^ + *R*^∗^(*β* + *ρ*_0_) = _1_ holds, hence, if *R*_1_, *R*_2_ and *S*_1_, *S*_2_ are the solutions of the two quadratic equations, then, in order to fulfil the previous equality, the *S* and *R* solutions corresponding to each other must have opposite signs. From this we obtain that under the assumptions of the proposition, there cannot exist a solution of (4) with two positive coordinates.

By linearizing (2) around the equilibria *E*_0_ and *E*_*R*_, respectively, and calculating the eigenvalues of the Jacobians of the linearized systems, we obtain the following results on the local stability properties of these two equilibria.

#### Proposition 2.5.

i. *E*_0_ *is locally asymptotically stable if and only if ℱ*_1_ *<* 0 *and ℱ*_2_ *<* 0.
ii. *E*_*R*_ *exists if and only if ℱ*_2_ *>* 0 *and it is locally asymptotically stable if and only if ℱ*_2_ *>* 0 *and ℱ*_3_ *<* 0.

### 2.2 Global dynamics

In this section, we turn to the study of the global dynamics of equation (2). To this end, we apply the Bendixson–Dulac criterion with choosing *D*(*S, R*) = 1*/*(*SR*) as a Dulac function. With this choice we obtain

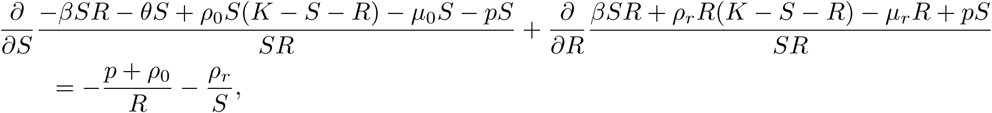

which is negative in the positive quadrant. Hence, using the Bendixson–Dulac theorem, we obtain that there is no periodic solution of (2). Applying the Poincaré–Bendixson theorem, it follows that all solutions tend to one of the equilibria. Based on this result and the local stability properties of the equilibria, we can give a complete characterization of the dynamics of system (2), depending on the threshold parameters *ℱ*_1_, *ℱ*_2_, *ℱ*_3_. To state our main result describing the global dynamics of the system, we introduce the notation

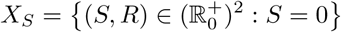

for the extinction space of sensitive cells.

#### Theorem 2.6.

*The global dynamics of equation* (2) *is completely determined by the threshold parameters ℱ*_1_, *ℱ*_2_, *ℱ*_3_ *as follows*.

i. *If ℱ*_1_ *<* 0 *and ℱ*_2_ *<* 0 *then the only equilibrium E*_0_ *is globally asymptotically stable*.
ii. *If ℱ*_2_ *>* 0 *and ℱ*_3_ *<* 0 *then E*_0_ *is unstable. The boundary equilibrium E*_*R*_ *is globally asymptotically stable. There is no coexistence equilibrium in this case*.
iii. *If ℱ*_1_ *>* 0, *ℱ*_2_ *<* 0 *and ℱ*_3_ *>* 0 *then E*_0_ *is unstable. The coexistence equilibrium is globally asymptotically stable on* 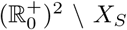. *There is no boundary equilibrium and E*_0_ *is globally asymptotically stable on X*_*S*_.
iv. *If ℱ*_1_ *>* 0, *ℱ*_2_ *>* 0 *and ℱ*_3_ *>* 0, *then E*_0_ *and E*_*R*_ *are unstable and the coexistence equilibrium E*_*C*_ *is globally asymptotically stable on* 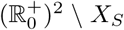. *The boundary equilibrium E*_*R*_ *is globally asymptotically stable on X*_*S*_.

The results of Theorem 2.6 are summarized in Table 1 where all possible combinations of the signs of the threshold parameters are listed along with the description of the existence and stability of the three possible equilibria. We note that the remaining two combinations of signs cannot be realized: the combinations *ℱ*_1_ *>* 0, *ℱ*_2_ *<* 0, *ℱ*_3_ *<* 0 and *ℱ*_1_ *<* 0, *ℱ*_2_ *>* 0, *ℱ*_3_ *>* 0 are excluded by Proposition 2.3. The regions defined by the signs of the three threshold parameters *ℱ*_1_, *ℱ*_2_, *ℱ*_3_ visualized in Figure 1. The ‘+’ and ‘−’ characters in the regions denote the signs of the parameters *ℱ*_1_, *ℱ*_2_, *ℱ*_3_. The figure was prepared with *β* = 0.0957, *ρ*_0_ = 0.1, *k* = 0.538, *µ*_0_ = 0.01, *µ*_*r*_ = 0.0262 and *p* = 0.01, while the parameters *θ* and *ρ*_*r*_ are varied between 0 and 0.1. Using Theorem 2.6, one can identify which of the equilibria is globally asymptotically stable on the given region. That is, *E*_0_ is globally asymptotically stable in the lower and middle regions on the right, *E*_*R*_ is globally asymptotically stable in the upper right and upper left regions, while *E*_*C*_ is globally asymptotically stable in the middle and lower regions on the left.

**Table 1:** Stability of equilibria depending on the threshold parameters.

| $\mathcal{F}_1$ | $\mathcal{F}_2$ | $\mathcal{F}_3$ | $E_0$ | $E_R$ | $E_C$ |
| --- | --- | --- | --- | --- | --- |
| − | − | ± | GAS | × | × |
| ± | + | − | unstable | GAS | × |
| + | − | + | unstable | × | GAS |
| + | + | + | unstable | unstable | GAS |

**Figure 1:**
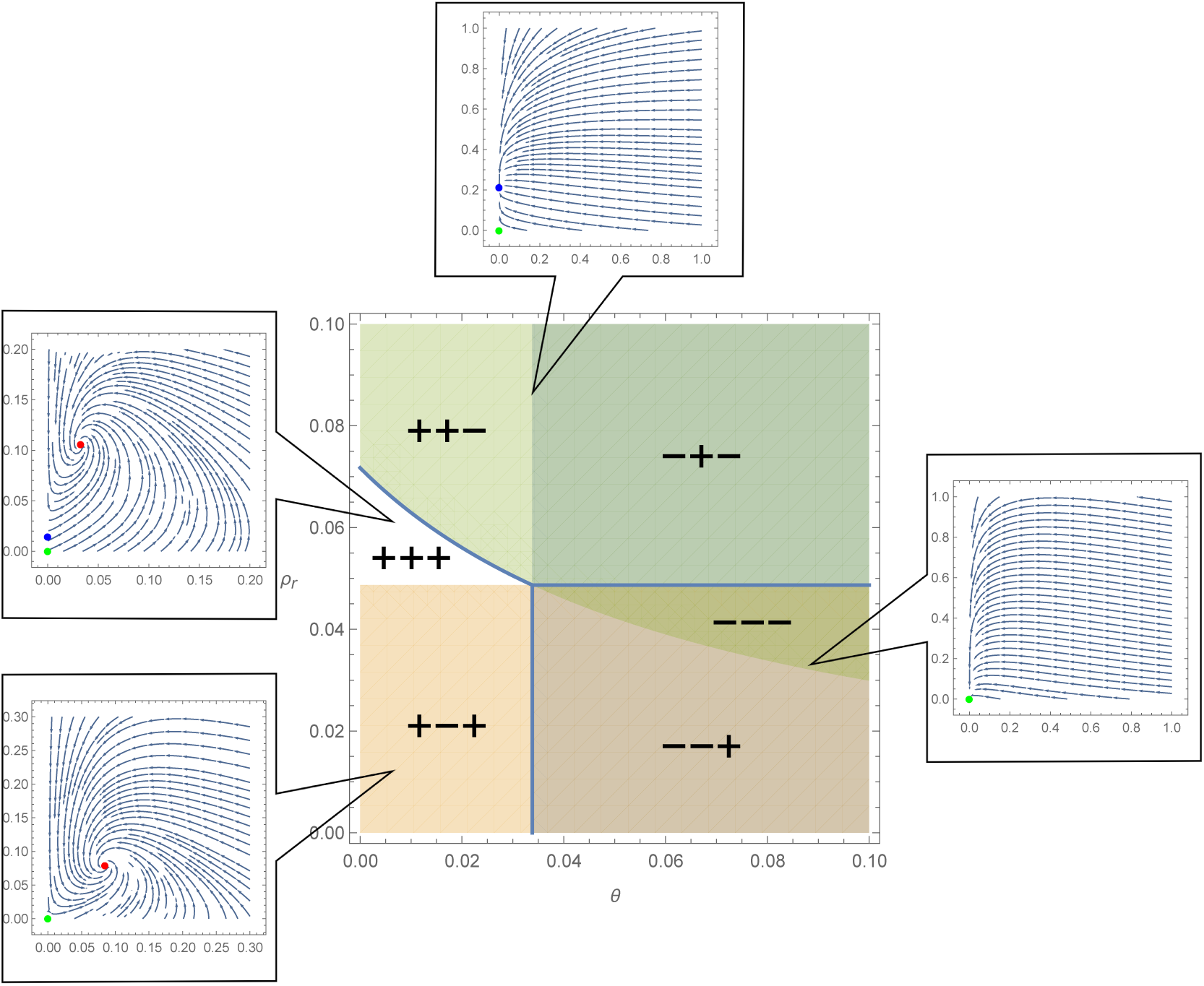
Regions corresponding to the different combinations of the signs of the three threshold parameters *ℱ*_1_, *ℱ*_2_, *ℱ*_3_.

## 3 Numerical simulations

### 3.1 The effect of drug concentration

We present numerical simulations to demonstrate what effects the change of the drug concentration *c* may induce. We study this through the change of the cytotoxic action induced cell mortality of sensitive cells *θ* and microvesicle production also depends on the concentration, assuming that these parameters are functions of the drug concentration, i.e. *θ* = *θ*(*c*) and *β* = *β*(*c*). We assume that both parameters are monotonically increasing in *c*, while the remaining parameters are fixed.

**Example 3.1**. We fix the parameter values

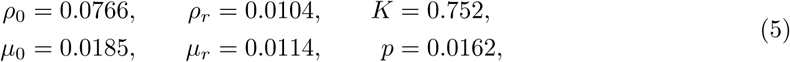

for which parameters the second threshold parameter has the fixed value *ℱ*_2_ = −0.00073 *<* 0. The level curves ℱ_1_ = 0 and ℱ_3_ = 0 divide the positive quadrant into three regions as shown in Figure 2. Using Theorem 2.6, we can identify the global dynamics of the system on all three regions: in the white region on the left, denoted by + − +, the coexistence equilibrium is globally asymptotically stable (except the extinction space of *R*, where *E*_0_ is globally asymptotically stable), while in the two remaining regions, *E*_0_ is globally asymptotically stable. Hence, when the curves leave the + − + region, a transcritical bifurcation occurs: the equilibrium *E*_*C*_ ceases to exist and *E*_0_ becomes globally asymptotically stable.

**Figure 2:**
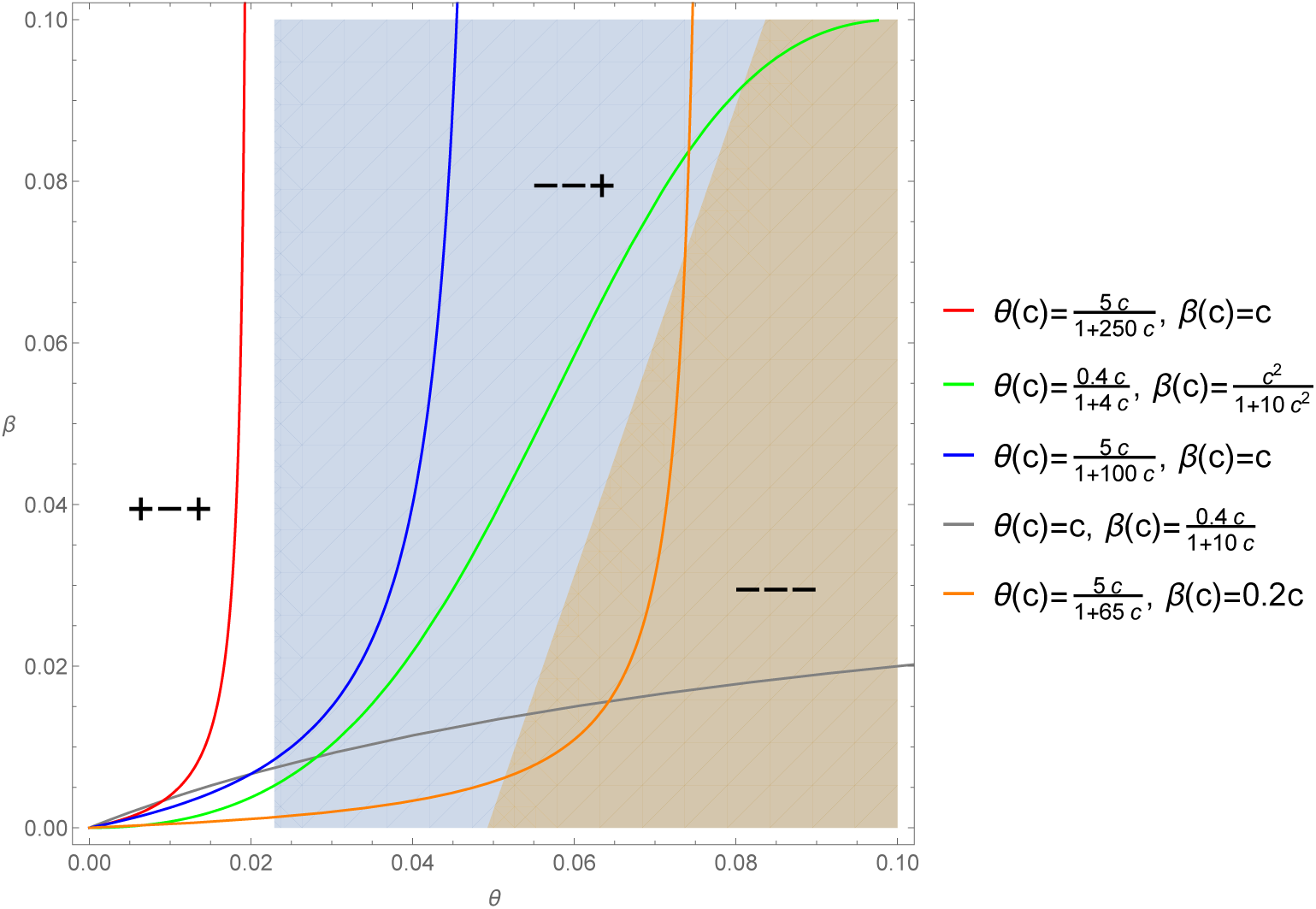
Possible scenarios for the global dynamics for different drug concentrations in the (*θ, β*)-plain for Example 3.1. The coloured curves represent possible transitions due to change in drug concentration, assuming different functional responses to the concentration.

Figure 4 (a) shows the total cancer mass, i.e. TCM = *S* + *R*, as a function of the drug concentration *c* for all five different functional forms of the increase of the concentration shown in Figure 2.

**Figure 3:**
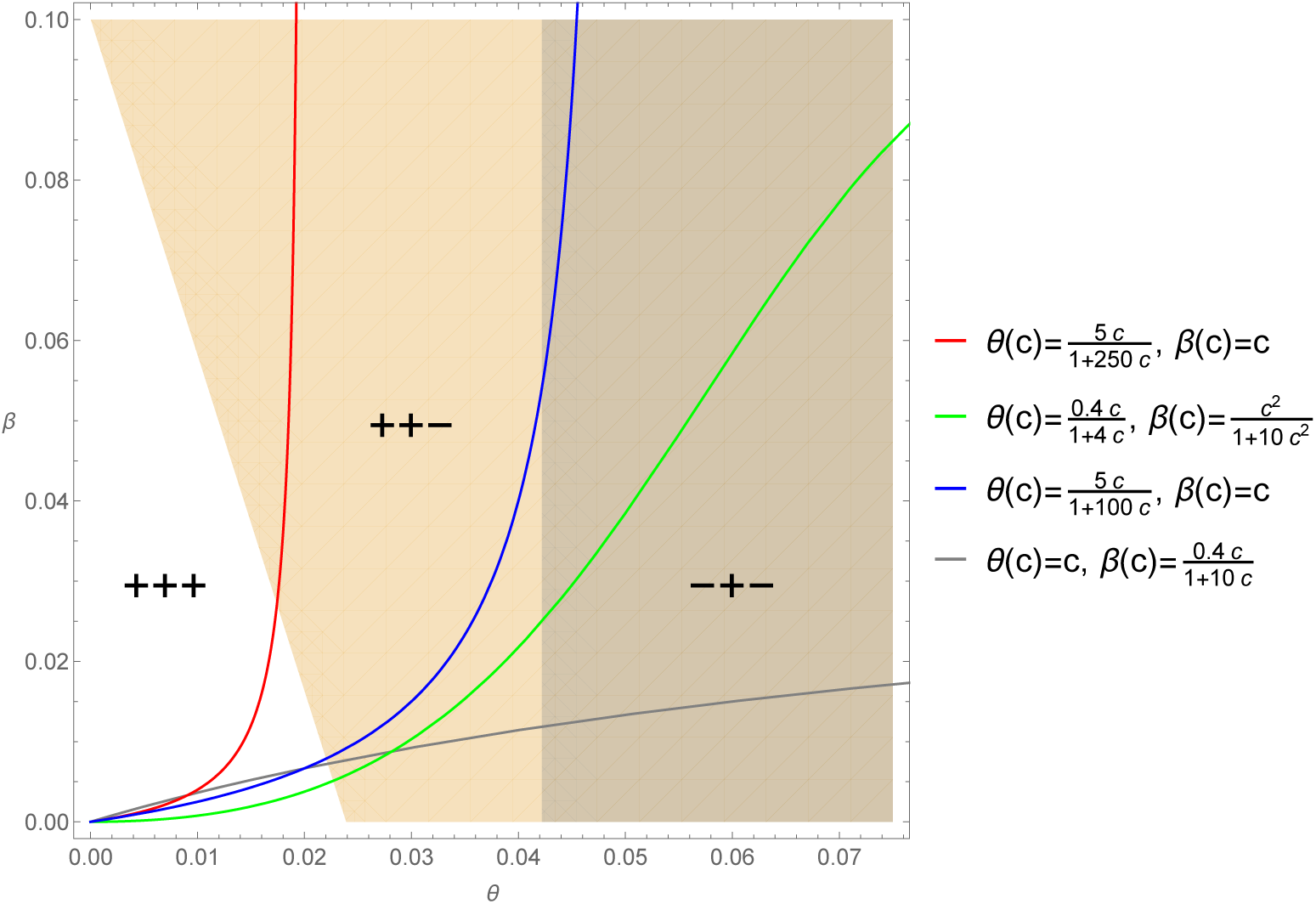
Possible scenarios for the global dynamics for different drug concentrations in the (*θ, β*)-plain for Example 3.2. The coloured curves represent possible transitions due to change in drug concentration, assuming different functional responses to the concentration.

**Figure 4:**
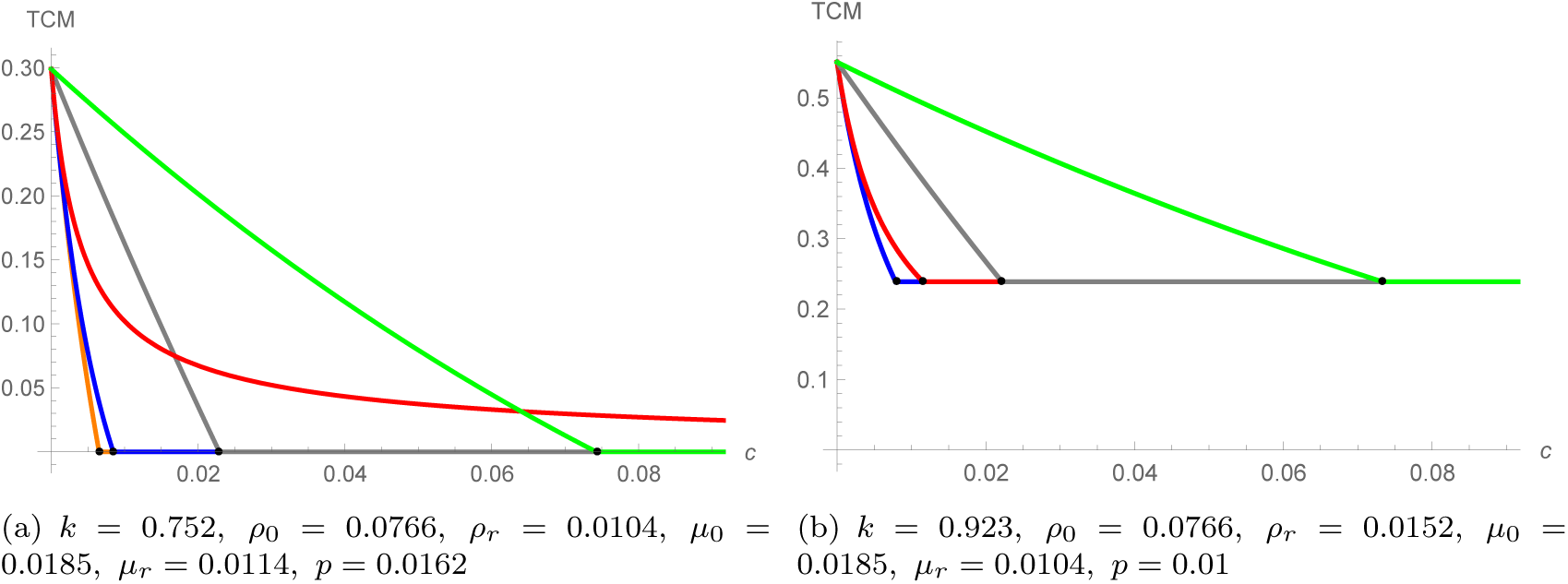
Total cancer mass (TCM) as a function of drug concentration *c* with different functional forms of *θ*(*c*) and *β*(*c*) in the case *ℱ*_2_ *>* 0. Functional forms are as given in Figures 2 and 3

**Example 3.2**. Let us now fix the parameter values as

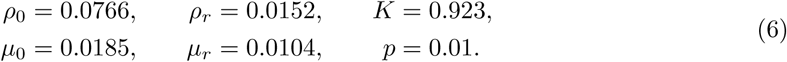

For these values, the threshold parameter ℱ_2_ has the fixed value ℱ_2_ = 0.036 *>* 0. For different values of the free parameters *β* and *θ*, we can experience different global dynamics depending on the sign of the remaining threshold parameters. Figure 3 shows the plain (*θ, β*), divided into three regions by the level curves ℱ_1_ = 0 and ℱ_3_ = 0. Using Theorem 2.6, we can determine which of the equilibria is globally asymptotically stable on that region.

Hence, one can see that *E*_*C*_ is globally asymptotically stable in the left white region, denoted by + + +, and *E*_*SR*_ is globally asymptotically stable in the remaining two regions. Assuming different functional forms for the dependence of *β* and *θ* on the drug concentration, we can see four possible sequences of transitions among the different regions depicted in Figure 3. When a curve passes through the boundary of the white region, a transcritical bifurcation occurs. Upon entering the white region, the previously globally asymptotically equilibrium *E*_*R*_ loses its stability and the newly arising coexistence equilibrium *E*_*C*_ becomes globally asymptotically stable.

Similarly as in the previous example, in Figure 4 (b) we depict the total cancer mass, TCM = *S* + *R*, as a function of the drug concentration *c* for the four functional forms of the increase of the concentration shown in Figure 3.

### 3.2 The effect of different chemotherapy regimes

In Figures 6 (a)–(d), we show the amount of sensitive and resistant tumour cells as a function of time with different regimes of chemotherapy for various values of the parameters. We assume that the chemotherapy drug is given to the patients. For the sake of simplicity, here we neglect the drug uptake by the tumour cells. We assume that the drug is given to the patients in regular time intervals. Hence, the chemotherapy concentration is given by the impulsive differential equation

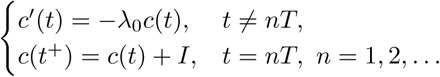

**Figure 5:**
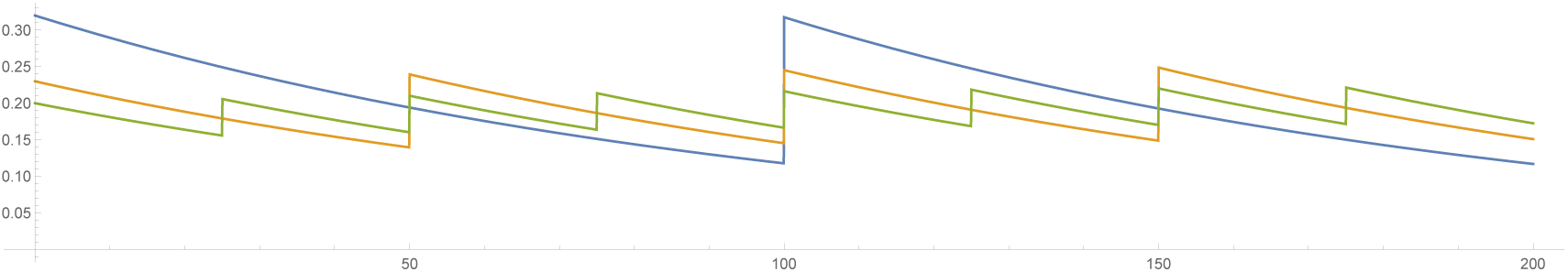
Drug concentration *c*(*t*) with three different regimes of drug dosage. We set *λ*_0_ = 0.01 for all three functions. The blue curve (longest interval between two treatments and largest drug amount) was prepared with parameter *I* = 0.2, the orange one (shorter period and lower drug amount) with *I* = 0.1, the green one (shortest period and lowest drug amount) with *I* = 0.05.

**Figure 6:**
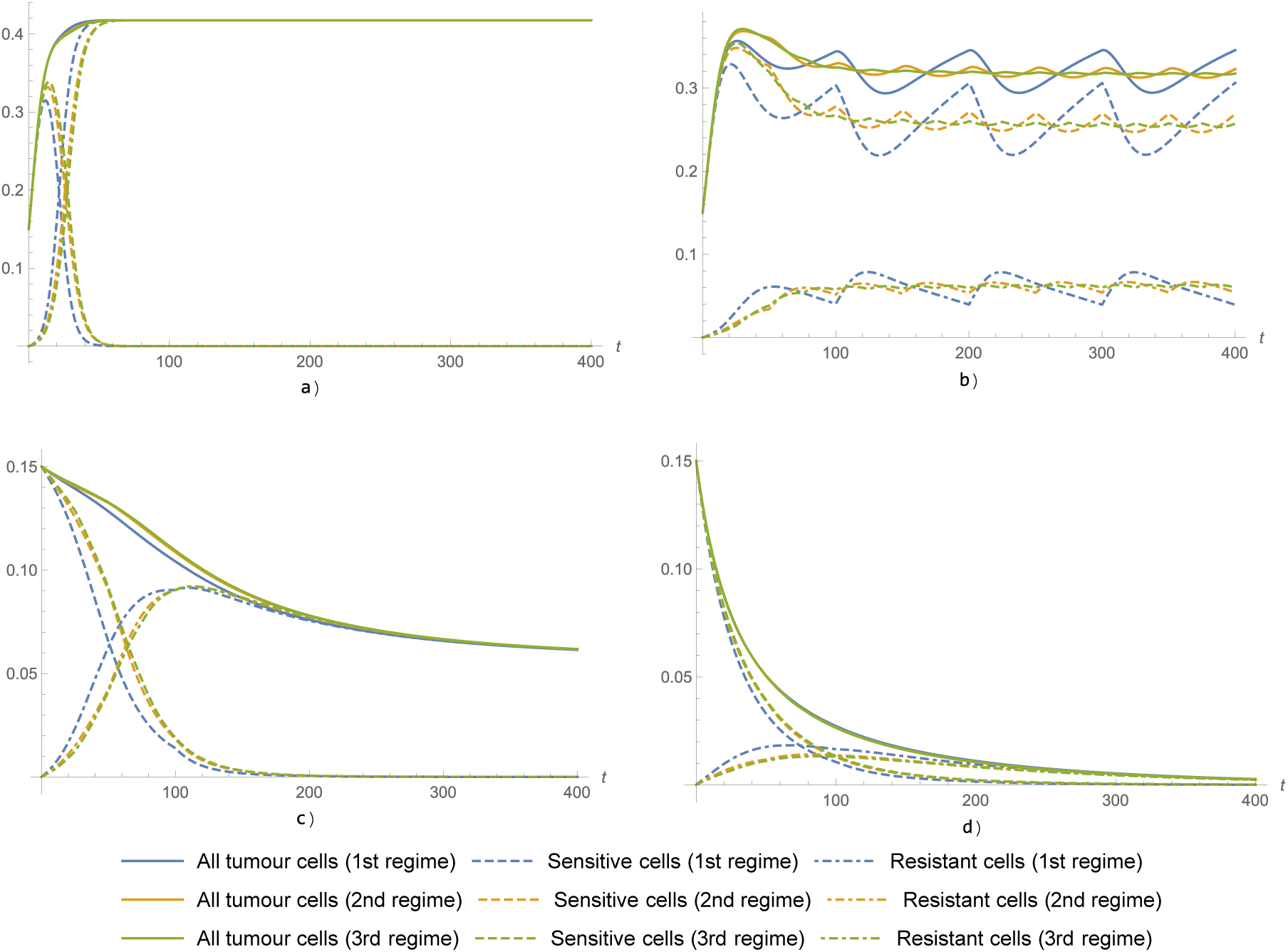
The number of all tumour cells, sensitive cells and resistant cells under the three different chemotherapy regimes shown in Figure 5 and various parameters. For all four simulations, *K* = 0.5, the rest of the parameters are as follows: a) *ρ*_0_ = 0.557; *ρ*_*r*_ = 0.281; *µ*_0_ = 0.0266; *µ*_*r*_ = 0.0409, b) *ρ*_0_ = 0.414; *ρ*_*r*_ = 0.154; *µ*_0_ = 0.0138; *µ*_*r*_ = 0.15, c) *ρ*_0_ = 0.157; *ρ*_*r*_ = 0.115; *µ*_0_ = 0.0257; *µ*_*r*_ = 0.0509, d) *ρ*_0_ = 0.166; *ρ*_*r*_ = 0.12; *µ*_0_ = 0.059; *µ*_*r*_ = 0.0662.

We compare three regimes which differ in the length of the time interval between receiving two doses of drug and in the amount of drug given at one treatment. The drug concentration *c*(*t*) corresponding to the three three different regimes of drug dosageare shown in Figure 5.

For illustratory purposes, we assume the following functional forms for the three drug-dependent parameters:

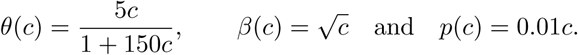

Figure 6 (a) can be interpreted as a failure of chemotherapy, as for all types of regimes, the tumour reaches a large size and all cells become resistant. Although there are slight differences between the regimes at the beginning of the therapy, finally, all three end in the same result. Figures 6 (b) and (c) show simulations where a partial success is achieved. However, in case (b), both types of cells are present, in contrary to case (c), where the sensitive cells tend to die out. In both cases, the regime with higher dose and longer period between two treatments seems to be the most effective as the total number of cells is the lowest in this case. Figure (d) shows a successful treatment, where the tumour disappears for all three types of regimes.

### 3.3 The effect of microvesicle transfer

An interesting question regarding the arising of chemotherapy resistance is in what extent the three ways of emergence contribute to resistance. In this subsection, we try to evaluate the effect of microvesicle transfer. Our numerical simulations suggest that microvesicle-mediated transfer might have an important role in determining the dynamics and the outcome of the treatment. It is not easy to provide a general description of this effect as it is clearly also heavily influenced by the rest of parameters, hence, here we only present two examples.

In Figure 7, we present a case where both a) without and b) with microvesicle-mediated transfer, sensitive cells die out and all cells become resistant. However, one may observe that the additional way of becoming resistant will increase the speed of the extinction of sensitive cells. In this case, the presence of microvesicle-mediated transfer does not affect the final size of the tumour.

**Figure 7:**
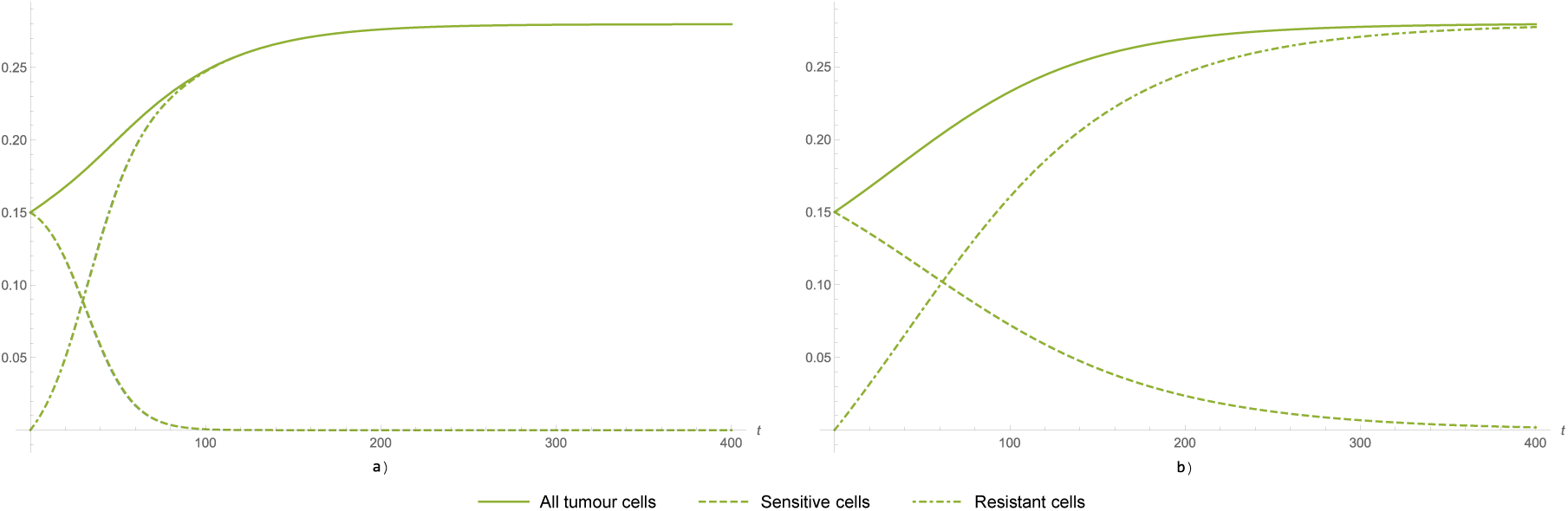
Dynamics a) with and b) without microvesicle-mediated transfer from sensitive to resistant cells. The parameters are set as *K* = 0.724, *ρ*_0_ = 0.0632; *ρ*_*r*_ = 0.0824; *µ*_0_ = 0.02; *µ*_*r*_ = 0.0366. The drug concentration dependent parameters are given as *β*(*c*) = 50*c/*(1+150*c*), *θ*(*c*) = 0.0108+0.0001*c* and *p*(*c*) = 0.01 + 0.0001*c*.

Figure 8 shows a situation which might seem paradoxic at first sight, as it shows that without microvesicle transfer, the total cancer mass becomes larger, although we have excluded one way of emergence of resistance. Hence, one would expect that if there is less possibility for the arising of resistance, the chemotherapy treatment might remain more efficient. However, one can observe that in this example, reproduction rate of sensitive cells is significantly higher than that of resistant cells, while death rate is significantly higher for the latter type of cells, i.e. by ignoring this way of cells becoming resistant, we allow a larger number of much more viable sensitive cells.

**Figure 8:**
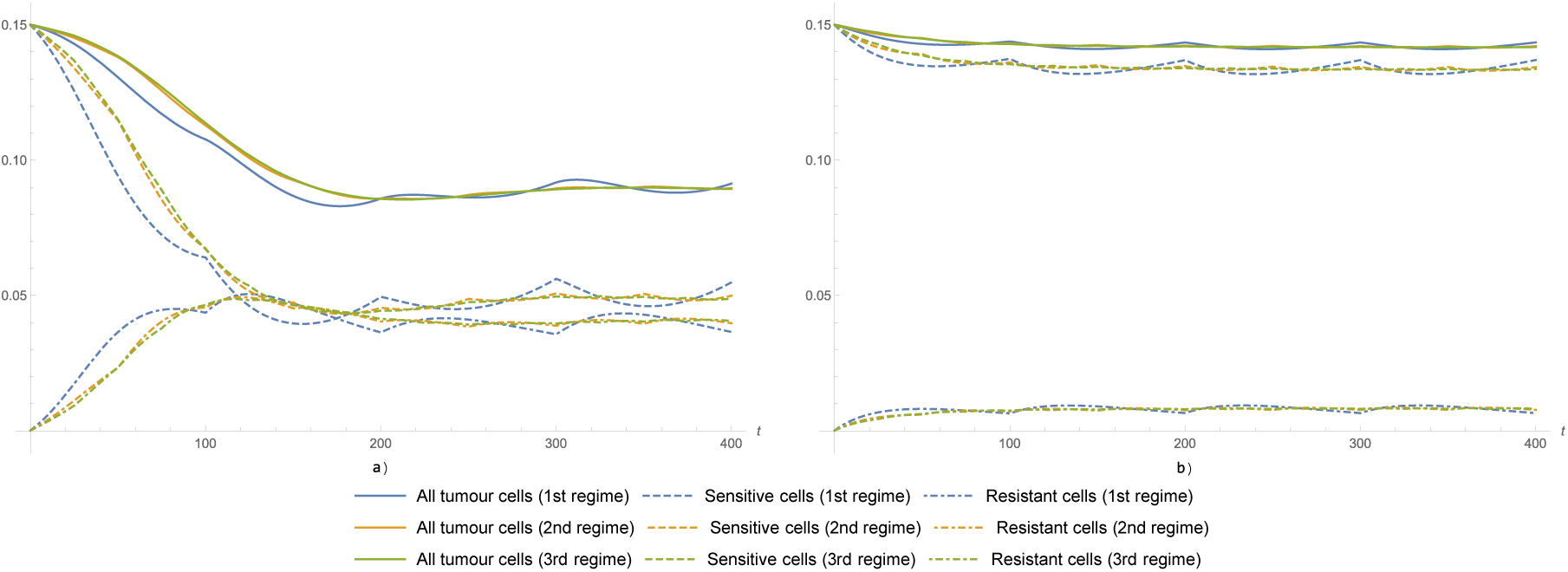
Dynamics a) with and b) without microvesicle-mediated transfer from sensitive to resistant cells. The parameters are set as *K* = 0.5, *ρ*_0_ = 0.3441; *ρ*_*r*_ = 0.157; *µ*_0_ = 0.089; *µ*_*r*_ = 0.0885.

## 4 Discussion

We established a mathematical model describing the evolution of tumour cells sensitive or resistant to chemotherapy. In the model, we considered three ways of emergence of chemotherapy resistance: as a result of the therapeutic drug: Darwinian selection, Lamarckian induction and, based on recent discoveries, the emergence of resistance via the transfer of microvesicles from resistant to sensitive cells, which happens in a similar way as the spread of an infectious agent. We calculated the possible equilibria of the system of differential equations describing the evolution of tumour cells and determined three threshold parameters which determine the global dynamics of the system. Using the Bendixson–Dulac theorem and the Poincaré–Bendixson theorem, we gave a complete description of the global dynamics characterized by the three threshold parameters. We demonstrated the possible effects of increasing drug concentration, and characterized the possible bifurcation sequences. We showed that increasing the drug-dependent parameters *β* and *θ* might turn the coexistence equilibrium *E*_*C*_ unstable and, depending on the rest of the parameters, either the resistant-only equilibrium *E*_*R*_ or the tumour-free equilibrium *E*_0_ becomes asymptotically stable. Another possible bifurcation is the appearance of the unstable equilibrium *E*_*R*_ while *E*_*C*_ remains asymptotically stable.

We note that in the presence of Lamarckian induction, no sensitive-only equilibrium exists, while in the absence of this phenomenon, such an equilibrium exists. Moreover, in the latter case, the existence of a coexistence equilibrium is only possible if a sensitive-only equilibrium exists as well (for details, see [5]).

In assessing the importance of development of resistance through microvesicle transfer, with the help of Table 1 and Figures 2–3, one can observe that that the presence of microvesicle transfer might turn a partially successful therapy to a failure, however, it is not able to ruin an otherwise successful therapy leading to extinction of the tumour. At the same time, as suggested by the simulations in Subsection 3.3, the absence of microvesicle transfer might even lead to an increase of total cancer mass if sensitive cells are much more viable than resistant cells.

## 5 Acknowledgements

A. Dénes was supported by the Hungarian National Research, Development and Innovation Office grant NKFIH PD 128363 and by the János Bolyai Research Scholarship of the Hungarian Academy of Sciences. G. Röst was supported by EFOP-3.6.1-16-2016-00008 and by the Hungarian National Research, Development and Innovation Office grant NKFIH KKP_129877 and 20391-3/2018/FEKUSTRAT.

## Notes

### Competing Interest Statement

The authors have declared no competing interest.

